# A robust example of collider bias in a genetic association study

**DOI:** 10.1101/028035

**Authors:** Felix R Day, Robert A Scott, Ken K. Ong, John R.B. Perry

## Abstract

A recent paper by Aschard *et al* described the potential for “collider bias” when adjusting for heritable covariates in genetic association studies^1^. However, in their examples the authors acknowledged that they could not exclude the possibility of a true biological explanation for the genetic association seen only in the adjusted model. Furthermore, the extent to which this bias could create a completely spurious genetic association, rather than just modify the magnitude of the effect,^2^ remains unclear.

Collider bias describes the artificial association created between two uncorrelated exposures *(A* and *B*) when a shared outcome (*X)* is included in the model as a covariate **(Figure 1).** We sought to definitively illustrate collider bias by deliberately inducing it to generate a biologically implausible SNP-phenotype association. Both sex (A) and autosomal genetic determinants for adult height (B) have causal effects on height (*X),* but are themselves implausibly correlated. We theorized that collider bias would induce false-positive associations between (only) autosomal height-associated genetic variants and sex when adjusting for height as a covariate.

**Figure 1.**
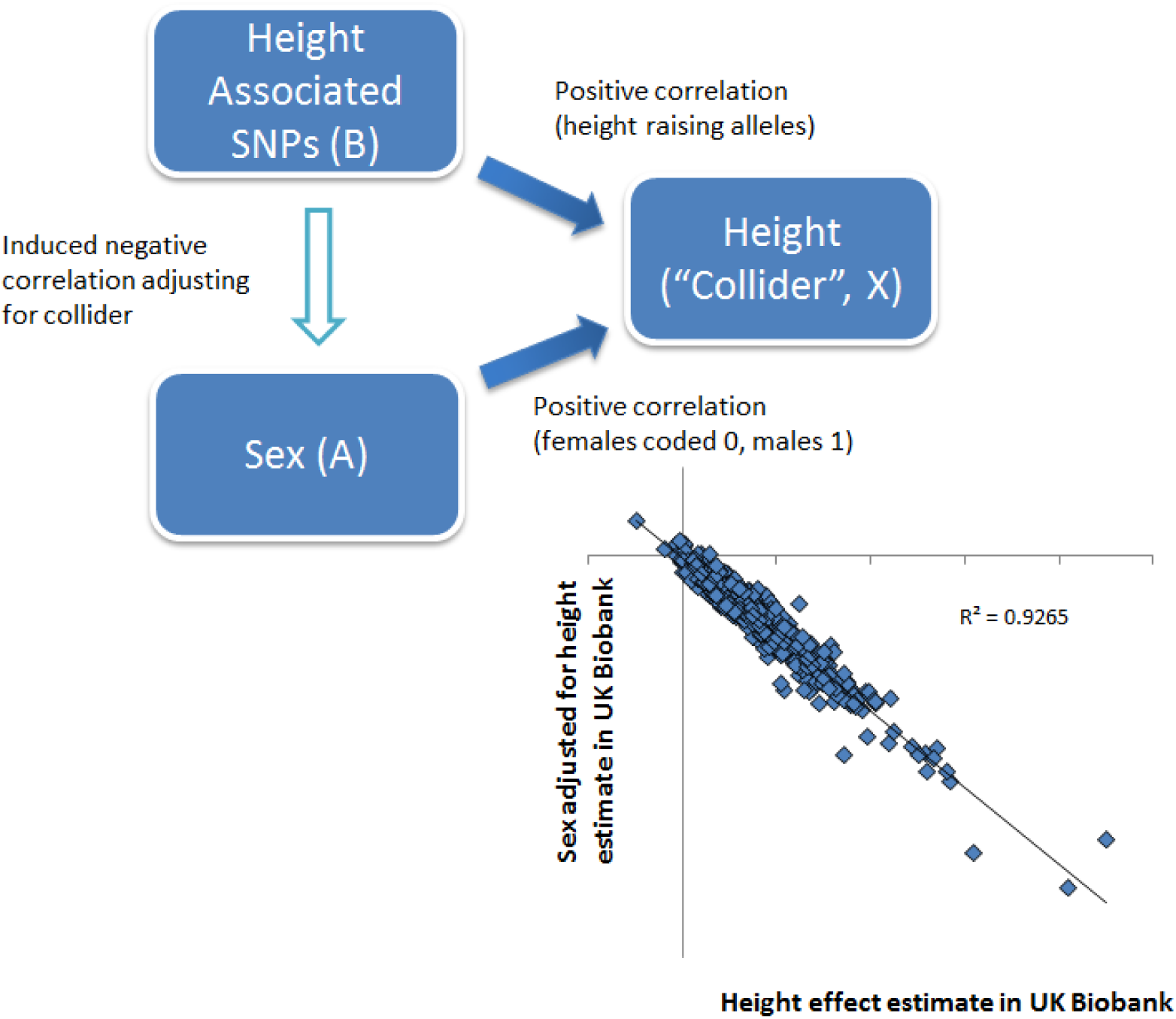
Spurious autosomal SNP effect estimates for sex, created by adjusting for height as a covariate, are almost perfectly correlated with SNP effect estimates for height. In this scenario of “collider bias”, adjustment for the “collider” height creates biologically implausible sex associations for the 697 previously identified genome-wide significant autosomal SNPs for height.

In a sample of 142,630 individuals of white European ancestry from the UK Biobank study^3^, we performed a genome-wide association study for sex using a linear mixed model and applying standard quality control metrics. As expected, in univariate models, no SNP reached genome-wide significance and test statistics for the 697 previously identified height SNPs^4^ conformed to a null distribution (P_min_=1.4×10^−3^, 27 SNPs P<0.05, ~35 by expected by chance). In contrast, when we repeated the analysis including height as a covariate, 222/697 height SNPs reached genome-wide significance for association with sex. Each height increasing allele exhibited the expected negative correlation with sex (i.e lower likelihood of being male), given the two causal exposures were aligned to be positively associated with height **(Figure 1).** This was exemplified by the three strongest signals in the reported GIANT height GWAS meta-analysis: *ZBTB38-*rs724016 (sex P_unadj_=0.05, P_adj_=7×10^−90^), *GDF5-*rsl43384 (P_unadj_=0.13, P_adj_=7×10^−71^) and *HMGA2-*rs8756 (P_unadj_=0.99, P_adj_=3×10^−34^), which all showed an apparent robust association with sex only in the height-adjusted model. Furthermore, amongst the 697 height-associated SNPs, their beta estimates for sex adjusted for height were almost perfectly correlated with their beta estimates for height adjusted for sex in this UK Biobank study sample **(Figure 1).**

In summary, we provide convincing evidence that adjusting for covariates can create apparently highly robust, but biologically spurious associations. The extent of this collider bias is almost perfectly inverse to the strength of the exposure-collider association. Consideration of causal inference modeling and unadjusted test statistics is therefore of great importance in the design and interpretation of genetic (and non-genetic) association studies.

**Note:** This article has been submitted as a “letter to the editor”. Further information regarding statistical methods is available upon request

## Acknowledgements

This work was conducted using the UK Biobank resource.

